# Establishment of a stable transgenic *g6pdM^1315-1443^* zebrafish line with glucose-6-phosphate dehydrogenase deficiency

**DOI:** 10.1101/2020.04.30.068981

**Authors:** Lu-Jun Shang, Jin Song, Hai-Xiong Xia, Yuan-Yuan Tuo, Ping-Ping Ren, Xi-Jun Wu, Yan-Hua Zhou, Jiao Jin, Chuan Ye, Zhi-Xu He, Li-Ping Shu

## Abstract

Glucose-6-phosphate dehydrogenase (G6PD) deficiency is the most common inherited enzymopathy in humans and is associated with a predisposition to hemolysis. However, there are few animal models to that adequately mimic associated human disease states that could be used to evaluate strategies to address clinical syndromes attributable to G6PD deficiency. In the present study, we aimed to establish a stable transgenic zebrafish model of G6PD deficiency that recapitulates the clinical manifestations of G6PD deficiency. We incorporated a stable transgene of G6PD lacking nucleotides from 1315 to 1443 denoted *Tg*(*zgata1:g6pd*^*M1315-1443*^-*egfp*). Functional analysis showed that *Tg*(*zgata1:g6pd*^*M1315-1443*^-*egfp*) transgenic zebrafish demonstrate a decrease in g6pd activity, reduced GSH levels and hemoglobin content, and increases in pericardial edema in response to α-naphthol exposure, similar to human subjects with G6PD deficiency. We detected no other significant phenotypic abnormalities compared to controls. Taken together, these observations indicate that the *Tg*(*zgata1:g6pd*^*M1315-1443*^-*egfp*) zebrafish line mirrors key clinical manifestations of G6PD deficiency in humans. This model may facilitate mechanistic studies and promote translational research related to G6PD deficiency.

## INTRODUCTION

Glucose-6-phosphate dehydrogenase (G6PD) deficiency is the most common inherited enzymopathy in humans, affecting about 400 million people worldwide [1]. The prevalence of G6PD deficiency geographically correlates with the distribution of malaria [2, 3] with a prevalence of 5-25% in persons residing in tropical Africa, the Middle East, tropical and subtropical Asia, some areas of the Mediterranean, and Papua New Guinea [1, 2]. In US, the prevalence of G6PD deficiency is about 10%, with black males being primarily affected [1]. In China, G6PD deficiency is concentrated in the South [4]. G6PD deficiency is polymorphic with over 400 mutations being identified [5]. In the Chinese population 30 mutations predominate with A95G, G392T, G487A, A493G, C592T, C1024T, C1360T, G1376T, and G1388A being the most common mutations. G6PD deficiency is usually asymptomatic but can manifest as hematonosis, including acute hemolytic anemia, chronic hemolytic anemia, and kernicterus upon exposure to certain pro-oxidative drugs or oxidative chemicals, including sulfonamide antibiotics, antimalarials, and fava beans [3]. Hemolysis is the most common symptom for G6PD deficient subjects mainly due to oxidative stress caused red blood cells (RBCs) destruction [6, 7]. G6PD deficiency mainly occurs in males because it is inherited in a manner of an X-linked recessive disorder, although women can also suffer from hemolysis due to G6PD deficiency [3, 8].

Human G6PD (hG6PD, EC:1.1.1.49) is encoded by the *G6PD* gene located in the distal long arm of the X chromosome at the Xq28 locus [9, 10] and contains 13 exons and 12 introns, spanning 18.5 kb [9, 10]. hG6PD consists of 515 amino acids and exists as a homodimer or tetramer (Fig. 1) [10, 11]. *G6PD* is highly polymorphic with over 400 variants, with most mutations resulting in a single amino acid substitution [5]. These mutations can lead to alterations in G6PD structure, including destabilization of tertiary complexes, leading to reduced biological activity. Functionally, G6PD is critical for the maintenance of redox homeostasis in cells, especially RBCs [12]. G6PD is the first and rate-limiting enzyme in the pentose phosphate pathway (PPP), and the sole means of oxidizing glucose-6-phosphate to 6-phosphogluconolactone, accompanied by the reduction of nicotinamide adenine dinucleotide phosphate (NADP^+^) to nicotinamide adenine dinucleotide phosphate (NADPH). NADPH maintains glutathione in its reduced form acting as a redox scavenger and mediator of redox homeostasis in RBCs [13]. PPP is the only source for NADPH in RBCs [12, 13]. As such, RBCs are more vulnerable to oxidative stress and associated damage than other cells. Accordingly, oxidative stress jeopardizes the biological functions of RBCs in G6PD deficient subjects due to low levels of NADPH and reduced GST, leading to loss of cell integrity with accompanying hemolysis.

**Fig. 1.**
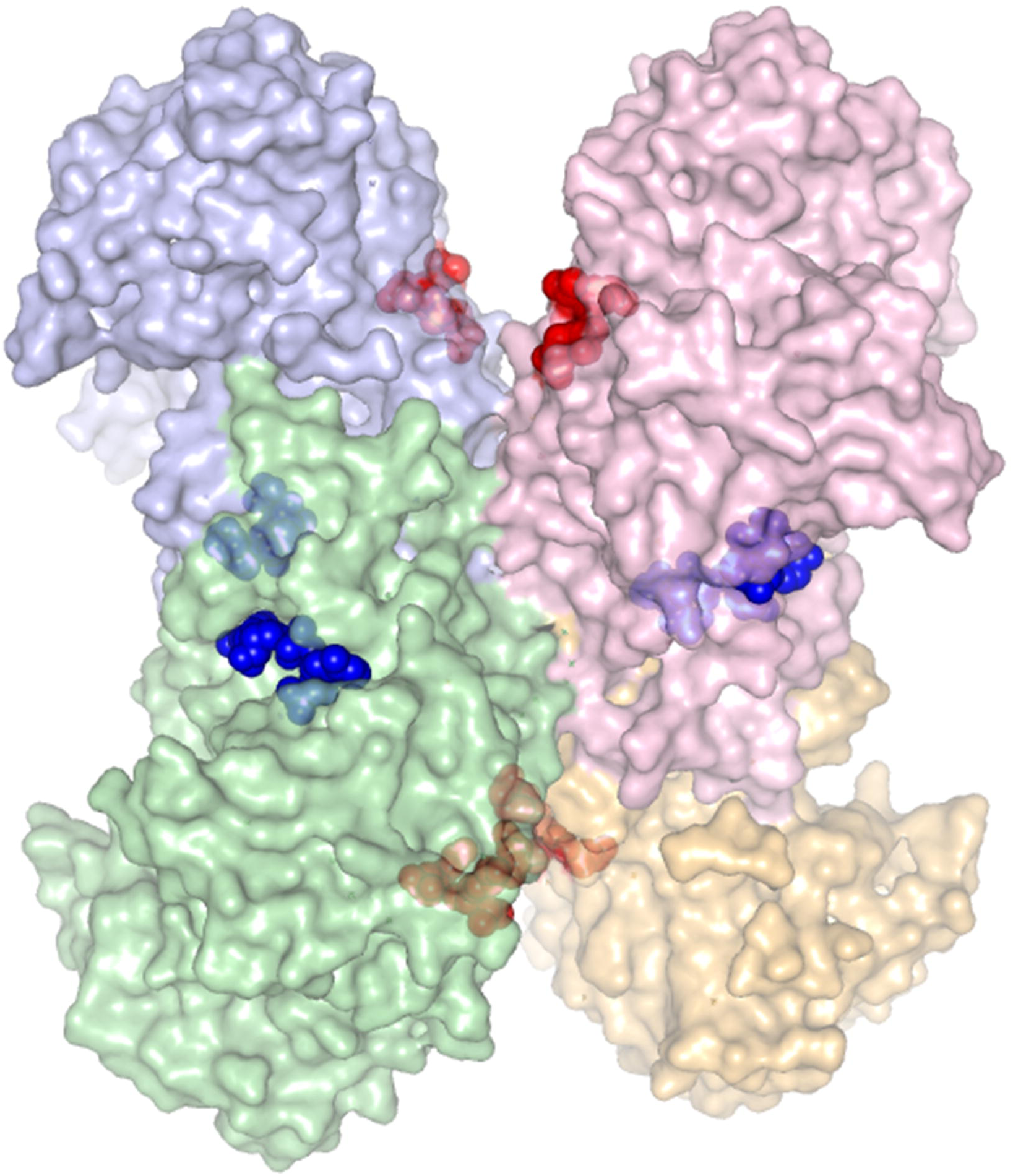
Human G6PD forms a tetramer (PDB ID: 2BHL). The location of most common destabilizing mutations, R454, R459, and R463, are highlighted in red and NADPH is blue.

Given the high prevalence of G6PD deficiency in humans and the vital roles of G6PD and redox metabolism in erythrocytic pathophysiology, development of appropriate research models mimicking human G6PD deficiency for mechanistic and functional studies that could lead to therapies that address G6PD deficiency is critical. Mouse models of G6PD deficiency have limited application because they demonstrate early embryonic death and lack a useful phenotype in the face of oxidative challenge [14]. Recently zebrafish models have increased in popularity as a model for hematonosis [15] because they allow for external genetic manipulations and provide a means of observing cellular events *in vivo* by virtue of transparent body compartments. Patrinostro et al. [16] established a zebrafish models of G6PD deficiency using morpholinos targeting the 5-prime exons of *g6pd*, but the model is limited to transient knockdown of *g6pd*. Here we report establishment of a stable transgenic *g6pd*^*M1315-1443*^ zebrafish line with a number of physiological features similar to human G6PD deficiency. This model of G6PD deficiency may enable research on the pathogenesis of hematonosis and translational research to address clinical conditions related to G6PD deficiency in humans.

## Results

### *zgata1:g6pd*^*M1315-1443*^-*egfp-pBSK-I-SceI* plasmid

C1360T, G1376T, and G1388A are three of the most common G6PD deficiency substitutions found in the Chinese population [17] and correspond to the mutations R454C, R459L and R463H, respectively (Fig. 1) [17]. These mutations are separate from the NADPH binding site and function by destabilizing the protein [17]. We developed a *g6pd* construct omitting the nucleotide sequence from 1315 to 1446 (Fig. 2A) to approximate the phenotype of these mutations. To this we fused egfp to enable fluorescence-based visualization *in vivo*, resulting in the construct *zgata1:g6pd*^*M1315-1443*^-*egfp-pBSK-I-SceI* which validated by electrophoresis and sequencing (Fig.S1A,B).

**Fig. 2.**
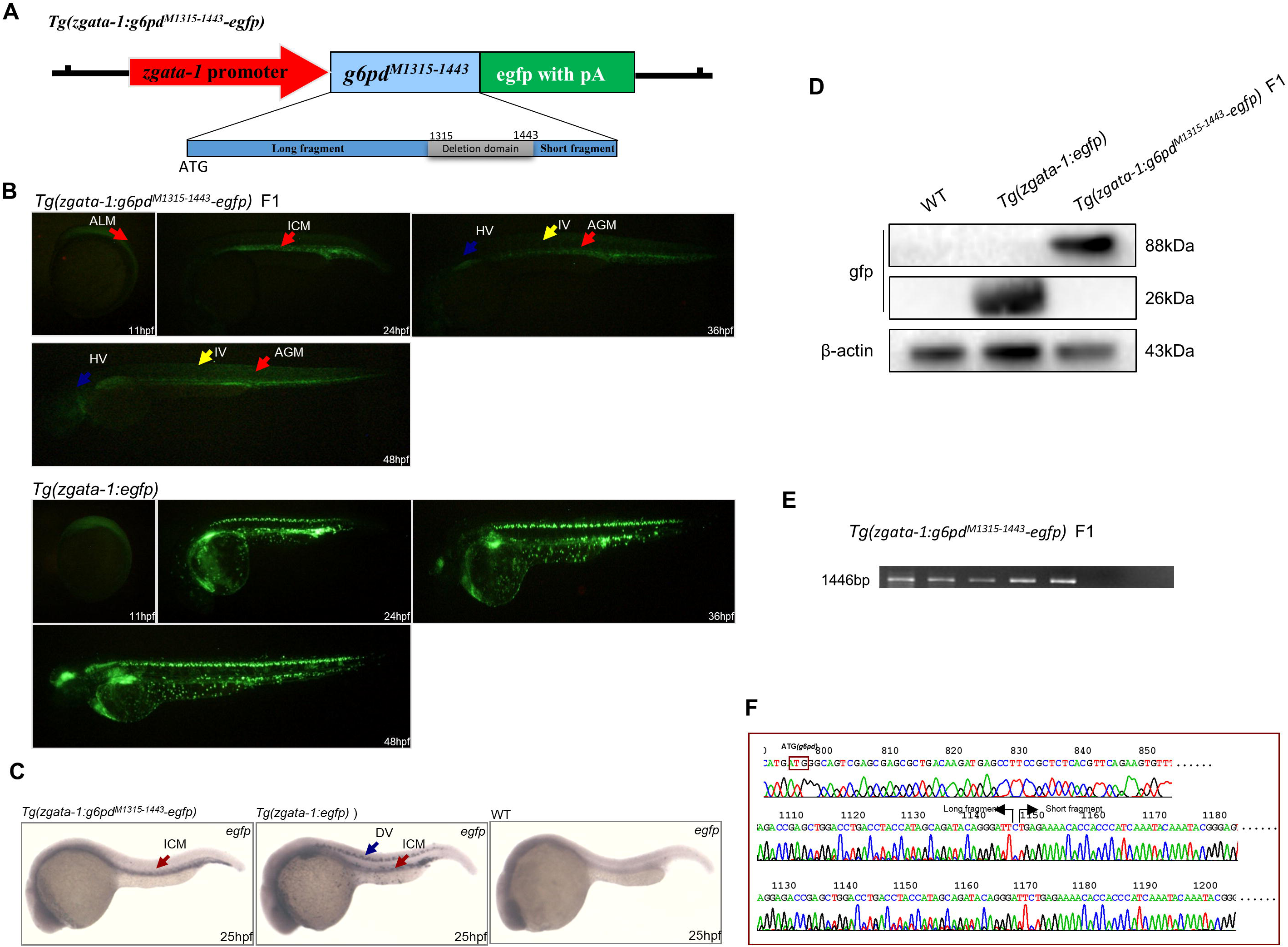
Expression of *Tg*(*zgata1:g6pd*^*M1315-1443*^-*egfp*) in the F1 generation of transgenic zebrafish. A: Structure of *Tg*(*zgata1:g6pd*^*M1315-1443*^-*egfp*) construct. B: Expression pattern of green fluorescence in F1 *Tg*(*zgata1:g6pd*^*M1315-1443*^-*egfp*) trangenic zebrafish and F1 *Tg*(*zgata1:egfp*) control trangenic zebrafish. HV: head vessels; IV: intersegmental vessels; AGM: aorta, gonads, and mesonephros. C: Expression of egfp mRNA was examined via *in situ* hybridization in F1 generation wildtype, *Tg*(*zgata1:egfp*), and *Tg*(*zgata1:g6pd*^*M1315-1443*^-*egfp*) zebrafish lines at 25hpf. DV: dorsal vessel; ICM: intermediate cell mass. D: Gfp protein expression was tested using Western blot in F1 generation wildtype (lane 1), *Tg*(*zgata1:egfp*) (lane 2), and *Tg*(*zgata1:g6pd*^*M1315-1443*^-*egfp*) (lane 3) zebrafish lines as measured by Western blot. E: PCR of genomic DNA shows presence of *g6pd*^*M1315-1443*^ in *Tg*(*zgata1:g6pd*^*M1315-1443*^-*egfp*) F1 generation of transgenic zebrafish. F: Sequence of fragments from panel E.

### Expression profile of *zgata1:g6pd*^*M1315-1443*^-*egfp-pBSK-I-SceI* plasmid

Expression of g6pd^M1315-1443^ in wildtype zebrafish was evaluated using fluorescence from egfp as a marker. *zgata1:g6pd*^*M1315-1443*^-*egfp-pBSK-I-SceI* was micro-injected into embryos at the one-cell stage (Fig.S2A), and expression of g6pd^M1315-1443^ was clearly evident by 9 hpf in the anterior lateral mesoderm and posterior lateral mesoderm (Fig.S2B). Egfp fluorescence was observed in the blood circulatory system at 24 hpf and decayed with time, vanishing at 48 hpf. No fluorescence was observed in wildtype zebrafish (Fig.S2B). In total, we injected 9,800 embryos with *zgata1:g6pd*^*M1315-1443*^-*egfp-pBSK-I-SceI* plasmid to obtain 5 F0 generation zebrafish with stable inheritance of egfp in the F1 generation. In the F1 generation, green fluorescence was initially observed at anterior lateral mesoderm at 11 hpf, overlapping the expression pattern of *gata1* (Fig. 2B). Egfp fluorescence was observed in the intermediate cell mass at 24 hpf (Fig. 2B). Subsequently, the fluorescence was observed in the head vessels (HV, blue arrow), intersegmental vessels (IV, yellow arrow), and aorta, gonads, and mesonephros (AGM, red arrow) at 36 and 48 hpf (Fig. 2B and Table 3). A more permissive pattern of egfp expression was observed in control zebrafish injected with an egfp control plasmid *Tg(zgata1:egfp)* (Fig. 2B).

**Table 3.** Comparison of expression levels in F1 *Tg(zgata-1:g6pd*^*M1315-1443*^-*egfp)* transgenic founders.

| Founder | Hematopoietic expression | Nonhematopoietic expression |
| --- | --- | --- |
| 1 | +<br>(ICM, AGM, HV, IV) | - |
| 2 | +<br>(ICM, AGM, HV, IV) | - |
| 3 | +<br>(ICM, AGM, HV, IV) | - |
| 4 | +<br>(ICM, AGM, HV, IV) | - |
| 5 | +<br>(ICM, AGM, HV, IV) | - |

Note: Expression levels were determined by observation at 24 hpf and 48 hpf using fluorescence microscope. +indicate relative expression levels; -: no observed. ICM: intermediate cell mass, AGM: aorta gonads and mesonephros, IV: intersegmental vessels, HV: head vessels.

### Identification the expression of *egfp* in F1 generation transgenic zebrafish line

To further validate the stable transgenic zebrafish line, we examined the expression of *efgp* mRNA in the F1 generation transgenic zebrafish via Whole-Mount in situ Hybridization (WISH) [18]. The assay was performed by using a digoxin labeled *egfp* anti-sense mRNA probe to examine the mRNA expression of *g6pd*^*M1315-1443*^ (Fig.S3). The probe was constructed from a linearized *egfp* product that was separated using electrophoresis and visualized after *BamHI* mediated digestion of the *egfp-pBSK* plasmid. The resultant product was subject to T3 RNA polymerase mediated transcription (Fig.S3). The *egfp* anti-sense mRNA probe was then used for *in situ* hybridization in wildtype, *Tg*(*zgata1:egfp*), and *Tg*(*zgata1:g6pd*^*M1315-1443*^-*egfp*) zebrafish lines at 25 hpf (Fig. 2C). The expression of *egfp* mRNA was detected at intermediate cell mass in both *Tg*(*zgata1:egfp*) and *Tg*(*zgata1:g6pd*^*M1315-1443*^-*egfp*) zebrafish lines and at dorsal vessel only in *Tg*(*zgata1:egfp*) zebrafish line. There was no *egfp* mRNA detected in wildtype zebrafish line (Fig. 2C). Moreover, egfp protein was detected in *Tg*(*zgata1:egfp*) and *Tg*(*zgata1:g6pd*^*M1315-1443*^-*egfp*) zebrafish lines at two molecular weights of 26 and 88 kDa, indicating egfp in the *Tg*(*zgata1:egfp*) transgenic zebrafish line and the g6pd^M1315-1443^-egfp fusion protein in *Tg*(*zgata1:g6pd*^*M1315-1443*^-*egfp*) transgenic zebrafish line respectively (Fig. 2D).

We next verified the genomic integration of *g6pd*^*M1315-1443*^ in F1 generation transgenic zebrafish line. The embryos for 5 F1 generation transgenic zebrafish were subject to genomic PCR to assay the expression of *g6pd*^*M1315-1443*^. The genomic DNA was isolated and fragmented as shown in Fig. 2E. There was a clear band with an approximate size of 1,550 bp, indicating the fragmentation of *g6pd*^*M1315-1443*^. The identity of the fragment was verified by Sanger sequencing as shown in the representative electrophorogram (Fig. 2F). Phenotypically and functionally the stable *Tg*(*zgata1:g6pd*^*M1315-1443*^-*egfp*) transgenic zebrafish line was otherwise indistinguishable from the parental strain or *Tg*(*zgata1:egfp*) in terms of acute hemolytic anemia, chronic hemolytic anemia, and kernicterus. To evaluate if we could elicit signs of G6PD deficiency using agents that cause hamatonoses in humans such as primaquine and α-naphthol, we next examined responses of the *Tg*(*zgata1:g6pd*^*M1315-1443*^-*egfp*) transgenic zebrafish line with a focus on g6pd activity, GSH levels, cardiovascular toxicity, and apoptosis of erythrocytes to primaquine and α-naphthol.

### G6pd activity in *Tg*(*zgata1:g6pd*^*M1315-1443*^-*egfp*) transgenic zebrafish line

Biochemical g6pd activity was first examined in F1 heterozygous and F3 homozygous *Tg*(*zgata1:g6pd*^*M1315-1443*^-*egfp*) transgenic lines. Of note, there were 4 F3 homozygous *Tg*(*zgata1:g6pd*^*M1315-1443*^-*egfp*) transgenic zebrafish lines generated (Table 4). g6pd^M1315-1443^-1 and g6pd^M1315-1443^-2 displayed global expression, but g6pd^M1315-1443^-3 and g6pd^M1315-1443^-4 exhibited specific tissue expression. Thusly, only g6pd^M1315-1443^-3 and g6pd^M1315-1443^-4 were used for the further assays. As shown Fig. 3A, the embryos for 4 F1 heterozygous *Tg*(*zgata1:g6pd*^*M1315-1443*^-*egfp*) transgenic zebrafish were assayed and there was no significant alteration in the g6pd activity at baseline as indicated by g6pd/6gpd ratio, compared to the wildtype zebrafish. Also, no significant change was observed in g6pd activity in embryos for 2 F3 homozygous *Tg*(*zgata1:g6pd*^*M1315-1443*^-*egfp*) transgenic zebrafish (g6pd^M1315-1443^-3 & g6pd^M1315-1443^-4) (Fig. 3B). However, g6pd activity was significantly decreased in embryos in the g6pd^M1315-1443^-4 transgenic zebrafish (F3) in the presence of 10 µM α-naphthol, evident from the marked reduction in the ratio of g6pd/6gpd, comparted to wildtype (Fig. 3C). Similar results were observed in the fluorescence spot test via the evaluation of fluorescence for reduced NADPH, indicating the significant decrease in g6pd activity in the g6pd^M1315-1443^-4 transgenic zebrafish (F3) exposed to 10 µM α-naphthol (Fig. 3D). Since g6pd activity is vital for the maintenance of the reducing power in cells [3], we subsequently examined the level of reduced GSH in the 2 F3 homozygous *Tg*(*zgata1:g6pd*^*M1315-1443*^-*egfp*) transgenic zebrafish (g6pd^M1315-1443^-3 & g6pd^M1315-1443^-4). In comparison to wildtype zebrafish, there was a significant decline in the level of reduced GSH that was observed only in g6pd^M1315-1443^-4 transgenic zebrafish (F3) embryos (Fig. 3E); and there was a dose-dependent reduction in reduced GSH level in g6pd^M1315-1443^-4 transgenic zebrafish (F3) embryos exposed to α-naphthol at 5, 10, and 20 µM (Fig. 3F). Nevertheless, we noted no phenotypic abnormalities in the two F3 homozygous *Tg*(*zgata1:g6pd*^*M1315-1443*^-*egfp*) transgenic zebrafish (g6pd^M1315-1443^-3 & g6pd^M1315-1443^-4) (Fig. 4A and B). In summary, these results show that our g6pd deficient zebrafish line exhibits similar biochemical features to human G6PD deficient subjects.

**Table 4.** Comparison of expression levels in F3 *Tg(zgata1-g6pd*^*M1315-1443*^-*egfp)* transgenic founders.

| Founder | Hematopoietic expression | Nonhematopoietic expression |
| --- | --- | --- |
| 1 | - | Global |
| 2 | - | Global |
| 3 | + | - |
| 4 | + | - |

Note: Expression levels were determined by observation at 18 hpf and 30 hpf using fluorescence microscope. + indicate relative expression levels; - no observed

**Fig. 3.**
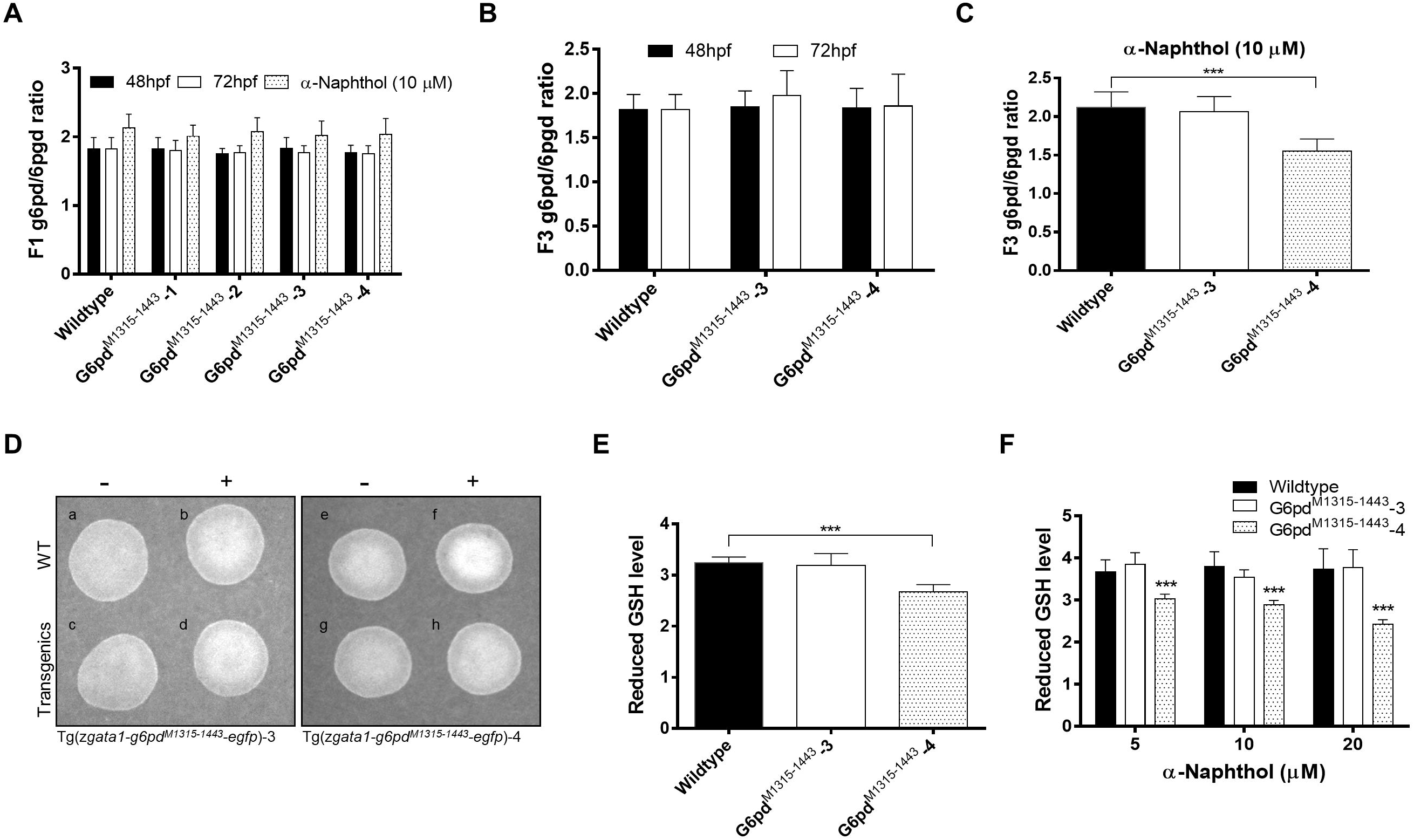
g6pd activity in *Tg*(*zgata1:g6pd*^*M1315-1443*^-*egfp*) transgenic zebrafish. A: g6pd activity in F1 heterozygous *Tg*(*zgata1:g6pd*^*M1315-1443*^-*egfp*) transgenic zebrafish in the presence and absence of α-naphthol (10 µM). B: g6pd activity in F3 homozygous *Tg*(*zgata1:g6pd*^*M1315-1443*^-*egfp*) transgenic zebrafish. C: g6pd activity in F3 homozygous *Tg*(*zgata1:g6pd*^*M1315-1443*^-*egfp*) transgenic zebrafish in the presence of α-naphthol (10 µM). D: Fluorescence spot test for g6pd activity in F3 homozygous *Tg*(*zgata1:g6pd*^*M1315-1443*^-*egfp*) transgenic zebrafish in the presence of α-naphthol (10 µM). Reduced GSH level in F3 homozygous *Tg*(*zgata1:g6pd*^*M1315-1443*^-*egfp*) transgenic zebrafish in the absence (E) or presence (F) of α-naphthol (10 µM).

**Fig. 4.**
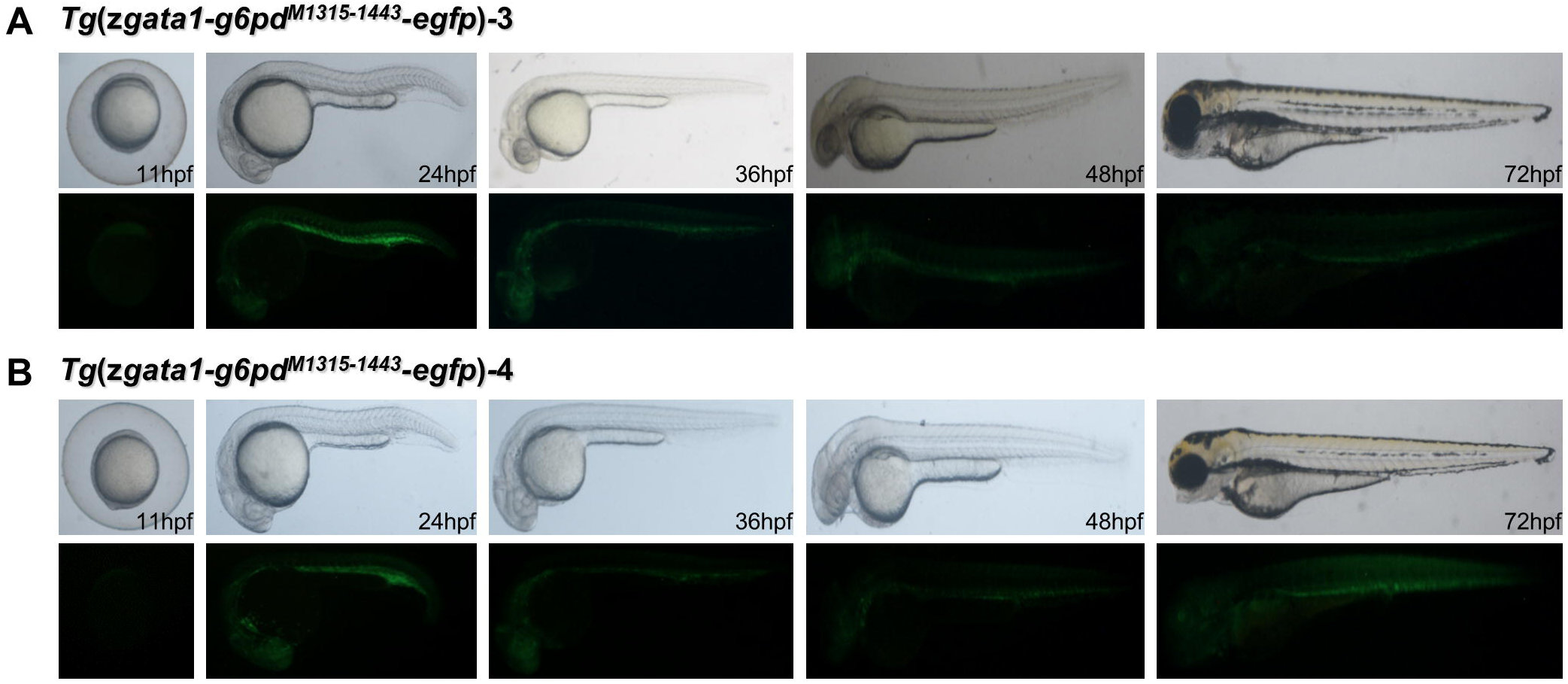
Phenotype of two F3 homozygous *Tg*(*zgata1:g6pd*^*M1315-1443*^-*egfp*) transgenic zebrafish lines (A: g6pd^M1315-1443^-3 & B: g6pd^M1315-1443^-4).

### Cardiovascular toxicity in F3 homozygous *Tg*(*zgata1:g6pd*^*M1315-1443*^-*egfp*) transgenic zebrafish line exposed to primaquine and α-naphthol

Prior zebrafish models of G6PD deficiency develop pericardial edema in response to oxidative challenge and resulting anemia [16]. To evaluate for heart malformations and pericardial edema secondary to G6PD deficiency, F3 homozygous *Tg*(*zgata1:g6pd*^*M1315-1443*^-*egfp*) transgenic zebrafish line were compared with wildtype. As shown in Fig. 5A and B, no morphological changes were found in the heart in two F3 homozygous *Tg*(*zgata1:g6pd*^*M1315-1443*^-*egfp*) transgenic zebrafish (g6pd^M1315-1443^-3 & g6pd^M1315-1443^-4) after exposure to 10 µM α-naphthol. However, we observed a remarkable degree of pericardial edema in g6pd^M1315-1443^-4 zebrafish upon exposure to 10 µM α-naphthol (Fig. 5C). Hemoglobin staining and erythrocyte staining showed a marked decrease in the level of hemoglobin in g6pd^M1315-1443^-4 zebrafish after exposure to 10 µM α-naphthol (Fig. 6A), which is correlated with the occurrence of pericardial edema. On the other hand, no abnormality was observed in erythrocyte phenotype (Fig. 6B). Furthermore, the apoptosis of erythrocyte was examined. The cells were sorted and those cells expressing egfp were collected for apoptosis assay. Compared to control, no significant difference in apoptosis was observed in erythrocytes derived from g6pd^M1315-1443^-4 zebrafish in the presence or absence of α-naphthol (Fig. 7).

**Fig. 5.**
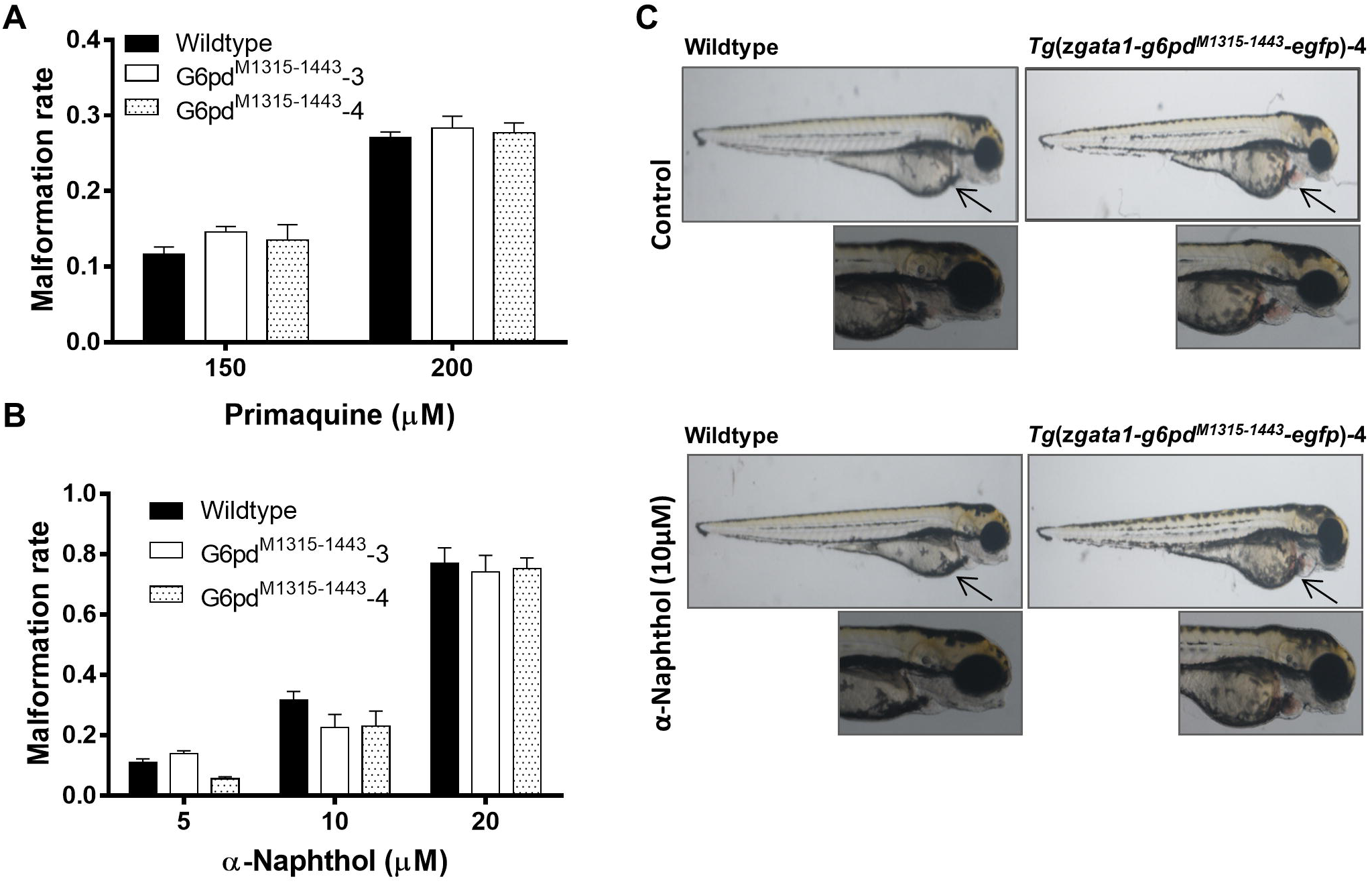
Cardiovascular toxicity-associated morphological abnormalities of F3 homozygous *Tg*(*zgata1:g6pd*^*M1315-1443*^-*egfp*) transgenic zebrafish line. A: Malformation rate of F3 homozygous *Tg*(*zgata1:g6pd*^*M1315-1443*^-*egfp*) transgenic zebrafish line in the presence of primaquine (A) and α-naphthol (B). Pericardial edema in g6pd^M1315-1443^-4 zebrafish in the absence (C) and presence (D) of α-naphthol.

**Fig. 6.**
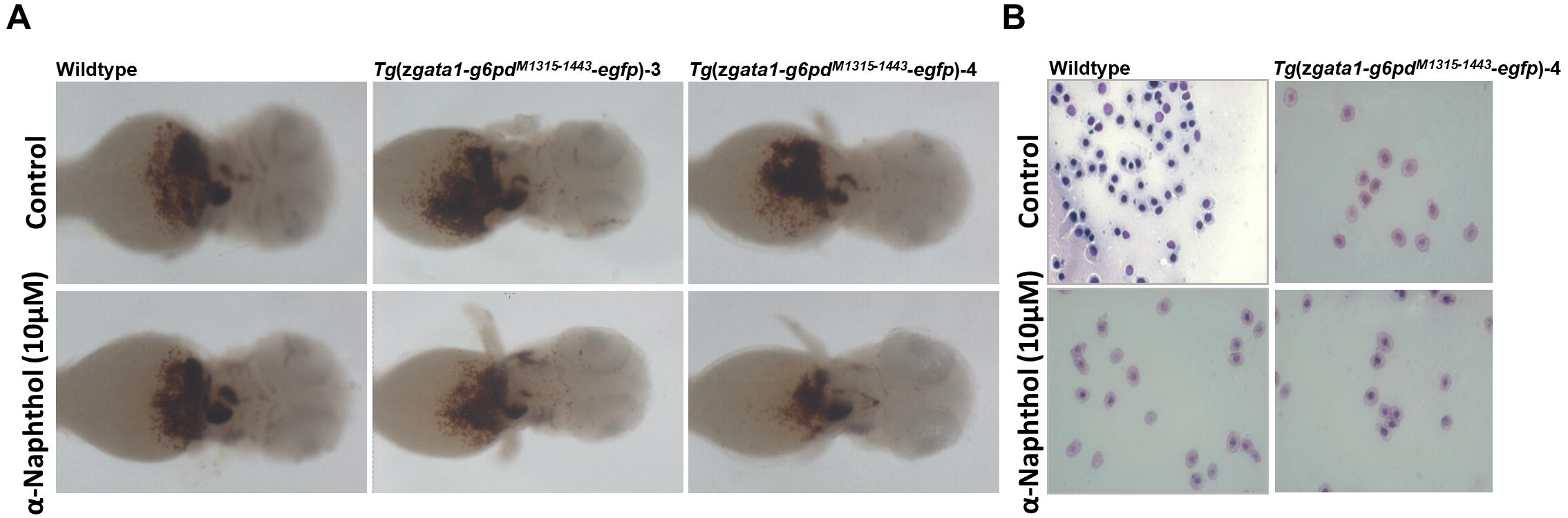
Hemoglobin staining and erythrocyte staining. A: Hemoglogin alteration in F3 homozygous *Tg*(*zgata1:g6pd*^*M1315-1443*^-*egfp*) transgenic zebrafish line in response to α-naphthol treatment. B: Erythrocyte phenotype in F3 homozygous *Tg*(*zgata1:g6pd*^*M1315-1443*^-*egfp*) transgenic zebrafish line in response to α-naphthol treatment.

**Fig. 7.**
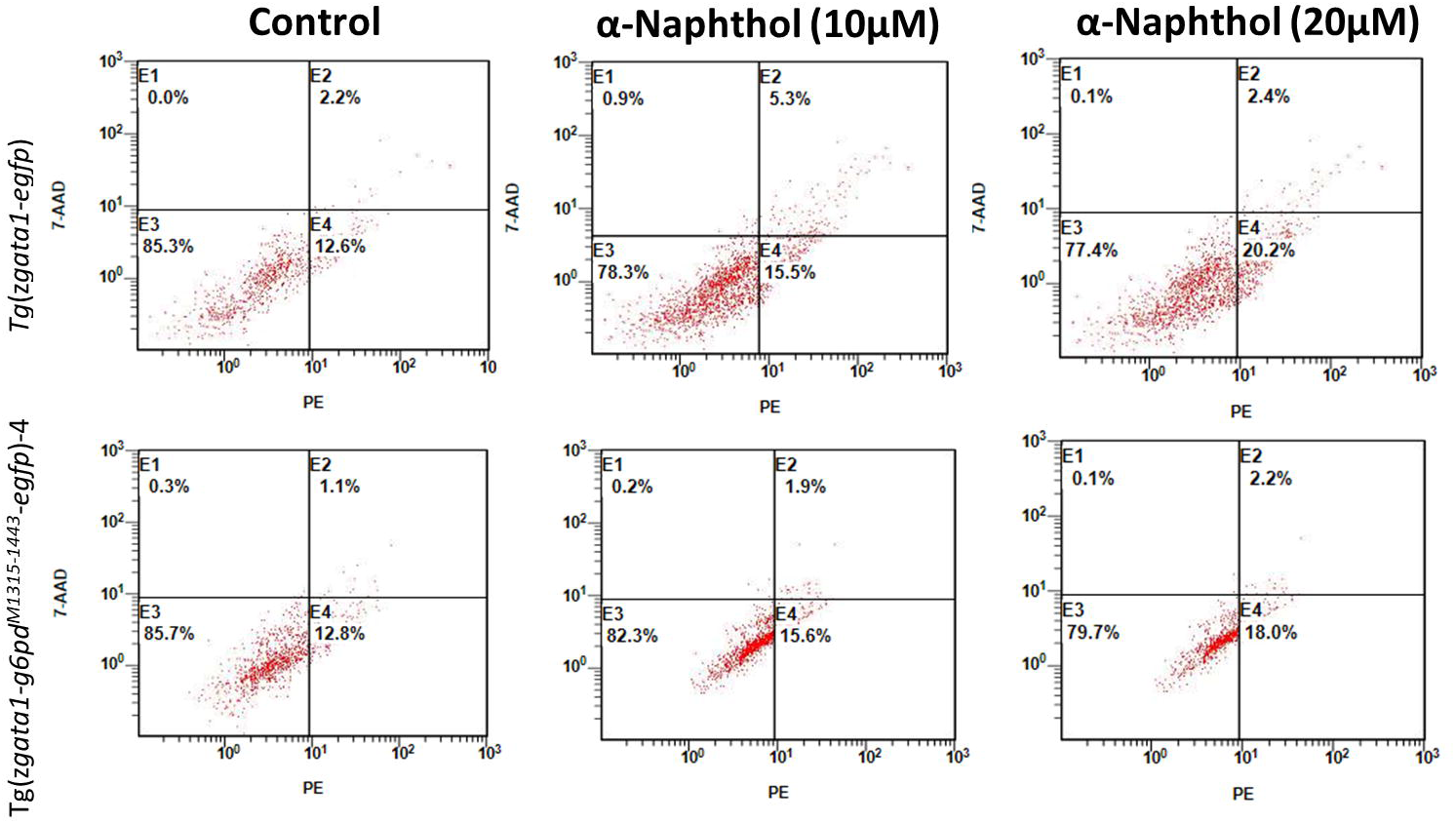
Apoptosis of erythrocyte in F3 homozygous *Tg*(*zgata1:g6pd*^*M1315-1443*^-*egfp*) transgenic zebrafish line with exposure to α-naphthol (10 and 20 µM).

## Discussion

A lack of suitable animal models stymies mechanistic and functional studies on G6PD deficiency and, consequently, hinders the translation to clinical practice. Here we report the development of a stable transgenic zebrafish line that mirrors the clinical manifestations of G6PD deficiency and provides the advantages of the zebrafish animal model. We showed that *Tg*(*zgata1:g6pd*^*M1315-1443*^-*egfp*) transgenic zebrafish demonstrated similar manifestations to human subjects with G6PD deficiency, with decreased g6pd enzymatic activity, reduced GSH level and hemoglobin content, but increased pericardial edema in response to α-naphthol exposure. The *Tg*(*zgata1:g6pd*^*M1315-1443*^-*egfp*) transgenic zebrafish had no other significant phenotypic abnormalities compared to controls. This stable transgenic animal model facilitates the investigation of the role of G6PD in erythrocytic pathophysiology and possesses great potential as a therapeutic target in the regulation of redox homeostasis, cell growth, and embryonic development.

G6PD deficiency is asymptomatic and the clinical manifestations occur in response to the exposure to certain drugs and chemicals [19]. It predisposes the subjects to hemolysis, resulting in great disease burden worldwide. Thus, it requires studies to in depth elucidate the underlying mechanism, so as to promote the translational research to curb with G6PD deficiency in clinic. In our model we used a deletion of nucleotides spanning bases 1315-1443 that corresponds to common disease alleles found in the Chinese population. These mutations are thought to affect the dimerization of G6PD, resulting in a compromised G6PD activity [17]. As such, a mutant g6pd was constructed and stably expressed in zebrafish with a nucleotide deletion from 1315 to 1443, in which contains C1360T, G1376T, and G1388A substations. Remarkably the *g6pd*^*M1315- 1443*^ transgenic zebrafish line did not show signs of G6PD deficiency at baseline, but did show the clinical features of G6PD when stimulated with chemical agents previously associated with G6PD deficiency phenotypes in humans. Compared to the previously reported models [16, 20], this model would facilitate further mechanistic and functional studies on G6PD deficiency, thanks to the transparent development of zebrafish allowing visualization of responses to stimuli and stable transgenes.

Mechanistically, the deletion of *g6pd* from 1315 to 1443 caused a miss of amino acid residues from 439 to 481. It would be expected that this deletion affect the dimerization of g6pd which is prerequisite for the activity of g6pd. The affected dimerization will lead to protein instability [17, 21]. Though we observed the expression of fused g6pd with egfp in *g6pd*^*M1315-1443*^ transgenic zebrafish, the relative low abundance may be attributed to the protein instability induced degradation. Further mechanistic studies are warranted to elucidate the effect of mutations on the function of g6pd.

Owing to the vital role of G6PD in the maintenance of NADPH pool and redox homeostasis, the activity of G6PD therefore would be singularly important to cells to cope with oxidative stress [13]. In RBCs, no other source generates NADPH than PPP, in which G6PD is the first and rate-limiting enzyme, justifying the singularly important role of G6PD in the biological events in RBCs. In contrast, cells with G6PD deficiency succumb to oxidative stress due to the inadequacy in the reductive power, reduced GSH, which highly hinges on the status of G6PD activity. As shown in F3 homozygous transgenic zebrafish without significant phenotypic change, g6pd activity was significantly compromised in the presence or absence of α-naphthol, compared to the control. In parallel with the decline in g6pd activity, there was a significant decrease in reduced GSH level in F3 homozygous transgenic zebrafish. Hemolysis is the most common consequence of G6PD deficiency in subjects exposed to prooxidants or antimalarial drug. Exposure to *g6pd*^*M1315-1443*^ transgenic zebrafish resulted in a decline in the content of haemoglobin, indicating the occurrence of hemolysis.

Taken together, our work presented a novel stable transgenic model of G6PD deficiency in zebrafish, which will facilitate the mechanistic and functional investigation in the role of G6PD in the maintenance of redox homeostasis, regulation of cellular signaling and embryonic development, erythrocytic pathophysiology. Also, utilization of this model would help predict the clinical hemolytic potential of drugs and promote translational research for infectious diseases (malaria) elimination.

## Methods

### Zebrafish strain and fish husbandry

Zebrafish lines were maintained in Zebrafish Core Facility, Guiyang Medical University using breeding and staging techniques previously described [15]. *Tg(zgata1:egfp)* zebrafish line was a gift from Dr. Min Deng (Institute for Nutritional Sciences, SIBS, Chinese Academy of Sciences, Shanghai, China). The protocol was approved by the Animal Experimentation Committee of Guiyang Medical University, Guizhou, People’s republic of China.

### zg6pd^*M1315-1443*^ primer design

Primers were designed to generate a deletion mutant zebrafish *g6pd* (*zg6pd*) lacking nucleotides 1315-1443, a sequence containing the most common variants seen in Chinese populations, G1376T and G1388A. <u>NCBI Reference Sequence: XM_694076.7</u> was used as a reference sequence. Primer sequences are listed in Table 1.

**Table 1.**
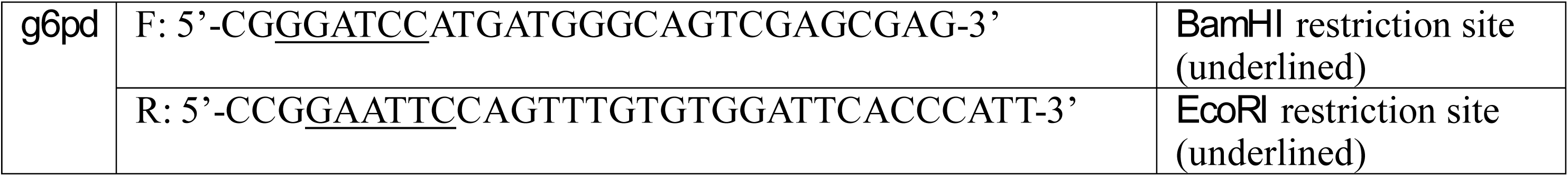
The sequences for the *g6pd* primers.

## Constructs

Recombinant plasmids were constructed as previously described by us [15]. In brief, total RNA was extracted from 10 zebrafish embryos at 24, 36, 48, 72, and 96 hpf using Trizol (Invitrogen, Carlsbad, CA, USA) and reverse transcribed into cDNA according to the manufacturer’s instructions (Fermentas, Canada). The resultant cDNA was used to synthesize *zg6pd*^*M1315-1443*^ using the primers listed in Table 1. *egfp*-*pCS*^*2+*^ and *egfp-pBSK* plasmids were constructed via *BamHI* and *EcoRI*-mediated digestion and T4 DNA ligase-mediated ligation. Then, the *zg6pd*^*M1315-1443*^ products and *egfp*-*pCS*^*2+*^ were digested by *BamHI* and *EcoRI* at respective restriction sites, and the recovered products were used to construct *zg6pd*^*M1315-1443*^-*egfp*-*pCS*^*2+*^ recombinant plasmid using T4 DNA ligase according to the manufacturer’s instructions (Fermentas, Canada). Similarly, *zgata1*:*pBSK*-*I-SceI* recombinant plasmid was constructed by *BamHI* and *EcoRI*-mediated digestion and T4 DNA ligase-mediated ligation. Subsequently, the *zgata1:g6pd*^*M1315-1443*^-*egfp*-*pBSK*-*I-SceI* recombinant plasmid was obtained from *zg6pd*^*M1315-1443*^-*egfp*-*pCS*^*2+*^ and *zgata1*:*pBSK*-*I-SceI* recombinant plasmids using *BamHI* and *EcoRI*-mediated digestion system and T4 DNA ligase-mediated ligation system. All recombinant plasmids (Table 2) were sequence verified. The *pCS*^*2+*^ and *pBSK* vectors and *pBSK* vector with *zgata1* promoter were kindly provided by Dr. Min Deng.

**Table 2.**
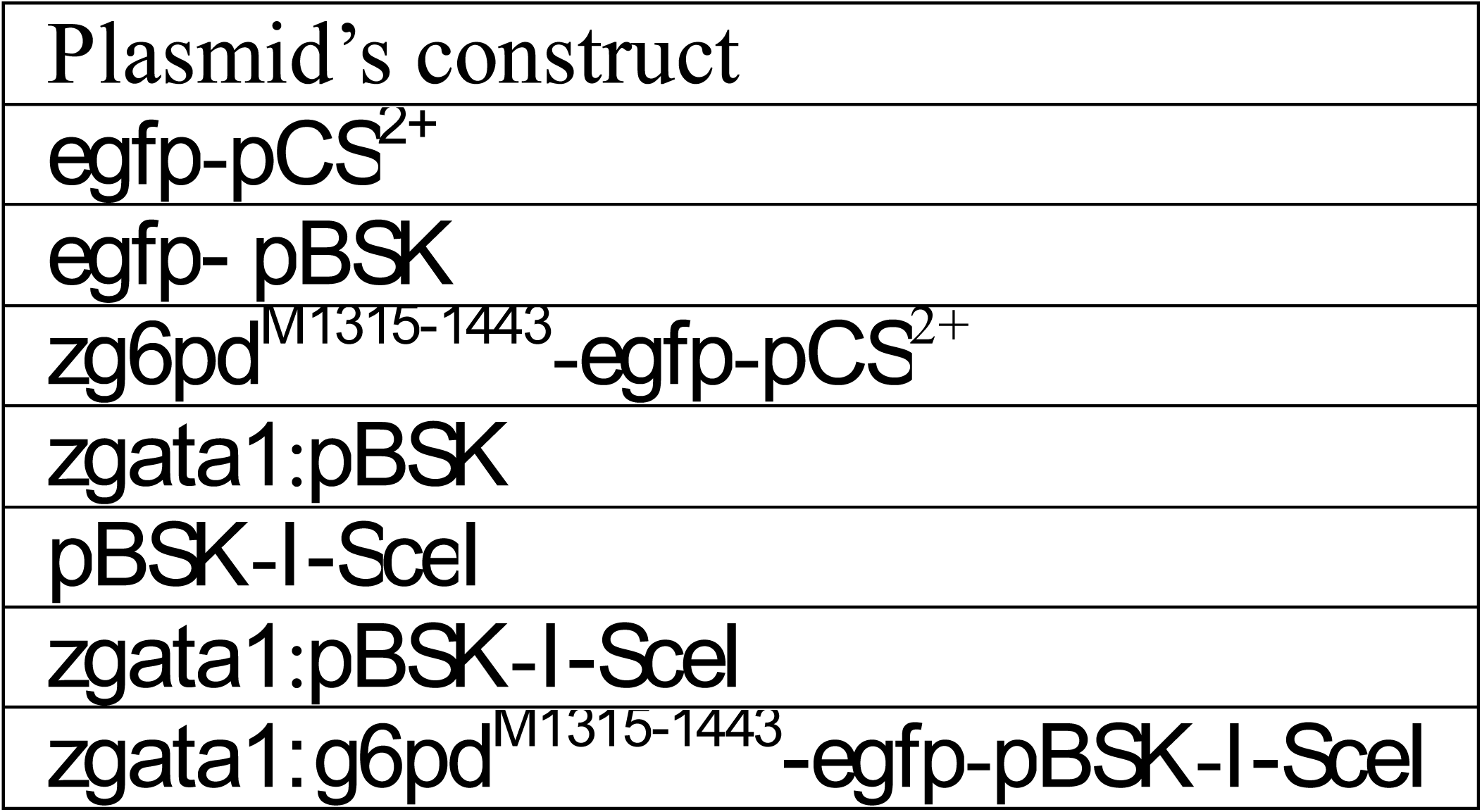
List of the plasmids used in this study.

### Micro-injection and generation of stable transgenic zebrafish lines

Plasmids were micro-injection to generate stable transgenic zebrafish lines as previously described with some modifications [15, 22]. In brief, plasmids were micro-injected into the developing cytoplasm of zebrafish embryos at the one-cell stage. The *I*-*SecI* meganuclease containing injection solution was prepared according to the manufacturer’s instruction (New England Biolabs, Ipswich, MA, USA). The potential founders (F0) were bred with wild-type fish and embryos were screened for green fluorescent protein (GFP) expression at 20–24 hpf. Those showing tissue-specific GFP expression were grown to adulthood (F1). The homozygous transgenic zebrafish line (F2) was descended from F1. Due to the possibility of mosaic integration in the germline, potential founders were only scored as non-transgenic after at least 100 of their offspring were shown to be negative for GFP expression.

### Microscopic imaging

To examine the expression of *g6pd*^*M1315-1443*^, the micro-injected embryos with *zgata1:g6pd*^*M1315- 1443*^-*egfp*-*pBSK*-*I-SceI* plasmids were held in egg water for imaging as previously described [15]. The embryos were screened for GFP expression using a Zeiss SteREO Discovery V12 fluorescence stereomicroscope. The images were acquired with the same microscope equipped with an AxioCam MRC5 digital camera and analyzed by AxioVision software.

### Western blotting analysis

Protein samples were collected from 30 embryos as previously described [15]. Briefly, the embryos were deyolked and homogenized. Harvested lysates were separated by SDS-PAGE. Primary antibodies against anti-egfp and anti-g6pd were obtained from Santa Cruz.

### PCR genotyping and DNA sequencing

Genomic DNA from 20 embryos of wild type, *zgata1:egfp*, and *zgata1:g6pd*^*M1315-1443*^-*egfp*-*pBSK*-*I-SceI* zebrafish lines were collected. Harvested DNA was used for PCR genotyping. PCR conditions were as follows: 94°C for 8 min; 35 cycles of 94°C for 40 sec; 65°C for 30 sec; 68°C for 90 sec; and then 68°C for 15 min. The resultant PCR samples were stored at −20°C for further DNA sequencing (Sunnybio, Shanghai, China). Genomic DNA from embryos of wildtype and *zgata1:egfp* zebrafish lines were used as control.

### *In situ* hybridization

A whole-mount mRNA *in situ* hybridization (WISH) assay was performed to examine the expression of *g6pd*^*M1315-1443*^ using digoxin labeled *egfp* probes. In brief, digoxin- and fluorescein-labeled *egfp* mRNA probes were transcribed from linear cDNA constructs according to the manufacturer’s instruction (Roche, Basel, Switzerland), and the WISH assays were performed as previously described [15]. Embryos were imaged with an AxioCam MRC5 digital camera and analyzed by AxioVision software. Embryos of wildtype and *zgata1:egfp* zebrafish lines were used as control.

### g6pd enzyme activity

Material from stable transgenic *zgata1:g6pd*^*M1315-1443*^-*egfp*-*pBSK*-Isce1zebrafish were used for functional assays. The activity of g6pd was evaluated as described previously with some modifications [23, 24]. In brief, g6pd activity was assayed in embryos when they were at 24 hpf and then exposed to α-naphthol (10 µM) over 48 hpf. After the treatment, the embryos were pelleted and snap frozen in liquid nitrogen and protein was extracted for the measurement of the ratio of G6PD over 6PGD using G6PD assay kit (ratio method) (Zhongshan Bio-Tech). The absorbance was measured at 650 nm. On the other hand, g6pd enzyme activity was assessed using fluorescence spot test as previously described [25, 26]. Briefly, the 24 hpf embryos were exposed to α-naphthol (10 µM) over 48 hpf and the g6pd enzymatic activity was measured based on the visual evaluation of fluorescence for reduced NADPH at 360-nm excitation and 460-nm emission. The fluorescence of NADPH is proportional to the activity of g6pd.

### GSH measurement

GSH levels were measured in the presence or absence of α-naphthol (5, 10, & 20 µM) over 48 hr using a reduced glutathione assay kit according to the manufacturer’s instructions (Nanjing Jiancheng Bioengineering Institute, Nanjing, China). In brief, the wildtype and transgenic embryos at 24 hpf were exposed to α-naphthol at 5, 10, and 20 µM for 48 hr and harvested at 72 hpf. After lysis and centrifugation, the supernatant was collected GSH detection in 96-well plates and the absorbance was measured at 405 nm.

### Cardiovascular toxicity evaluation

The cardiovascular toxicity was evaluated in wildtype and transgenic lines by assessment for pericardial malformation and pericardial edema in the F3 generation. F3 generation embryos were exposed to α-naphthol at 5, 10, and 20 µM for 48 hr and primaquine at 150 and 200 µM for 72 hr. Zebrafish were visualized and the images acquired at indicated phenotypic endpoints using a dissecting stereomicroscope.

### Staining and imaging (Hemoglobin staining & Erythrocyte staining)

O-dianisidine-mediated hemoglobin staining was performed as described previously [27]. Wildtype and F3 generation homozygous mutant embryos were exposed to 10 µM α-naphthol and harvested at 72 hpf. The dechorionated live embryos were stained with 0.6 mg/mL o-dianisidine and dehydrated through graded ethanol washes of 25%, 50%, and 100%. The processed embryos were imaged using a Nikon stereomicroscope with mounted NES imaging system. The images were analyzed using Image J. Erythrocyte staining was performed as previously described [28]. Wildtype and transgenic embryos at 24 hpf were exposed to 10 µM α-naphthol for 48 hr and harvested at 72 hpf. The erythrocytes were collected and stained with Wright-Giemsa or Prussian blue [29]. Images were obtained using NES system under Zess microscope.

### Flow cytometry

Apoptosis was detected using a PE annexin-V apoptosis detection kit according to the manufacturer’s instruction (BD, USA). Wildtype and transgenic homozygous embryos were exposed to α-naphthol at 5, 10, and 20 µM for 48 hr and harvested at 72 hpf. Erythrocytes were collected and stained with annexin-V and 7-AAD solution for 15 min in the dark. Subsequently, the cells samples were subject to flow cytometry (Beckman FC500)

### Statistical analysis

Data are expressed as mean ± SD. One-way analysis of variance (ANOVA) followed by Tukeys multiple comparison procedure was used for comparisons of multiple groups. P<0.05 was considered to be statistically significant. The assays were performed at least three times independently.

## Acknowledgement

This project was supported in part by the National Natural Science Foundation of China (31860325, 31360285), and in part by the Guizhou Province’s Science and Technology Major Project (Qian-P-Ren[2017]5611 & Qian-P-Ren[2019]5406), and in part by the Non-profit Central Research Institute Fund of Chinese Academy of Medical Sciences(NO.2018PT31048, 2019PT310013). We would like to thank Drs. Zhi-Wei Zhou and Kenneth Westover from UT Southwestern Medical Center, Dallas, US for their kind help on human G6PD structure analysis.

## Author contribution

Conceptualization: Li-Ping Shu and Zhi-Xu He; Methodology: Yan-Hua Zhou; Formal analysis: Yan-Hua Zhou, Hai-Xiong Xia, Yuan-Yuan Tuo, Lu-Jun Shang, Jin Song; Inverstigation: Yan-Hua Zhou, Hai-Xiong Xia, Yuan-Yuan Tuo, Lu-Jun Shang, and Jin Song; Resources: Li-Ping Shu, Zhi-Xu He and Chuan Ye; Writing: Zhi-Xu He, and Li-Ping Shu; Funding: Zhi-Xu He and Li-Ping Shu.

## Supporting data

Fig.S1. Construction of *zgata1:g6pd*^*M1315-1443*^-*egfp-pBSK-I-SceI* plasmid. A: The electrophoresis of *zgata1:g6pd*^*M1315-1443*^-*egfp-pBSK-I-SceI* plasmid fragmentations after BamHI, EcoRI, and XhoI mediated digestion. Lane 1: DNA marker, land 2: *zgata1:g6pd*^*M1315-1443*^-*egfp-pBSK-I-SceI* plasmid digestion products, lane 3: blank control. B: The sequence of *zgata1:g6pd*^*M1315-1443*^-*egfp-pBSK-I-SceI* plasmid.

Fig.S2. Expression of *zgata1:g6pd*^*M1315-1443*^-*egfp-pBSK-I-SceI* plasmid in wildtype zebrafish. A: Exhibition of of micro-injection of *zgata1:g6pd*^*M1315-1443*^-*egfp-pBSK-I-SceI* plasmid into embryo at the one-cell stage. B: Expression profile of g6pd in wildtype zebrafishe with or without injection of *zgata1:g6pd*^*M1315-1443*^-*egfp-pBSK-I-SceI* plasmid at 9, 24, 30, and 48 hpf.

Fig.S3. Synthesis of anti-sence egfp mRNA probe for the identification the expression of *egfp* in F1 generation transgenic zebrafish.

## Reference

1. Nkhoma, E.T., et al., The global prevalence of glucose-6-phosphate dehydrogenase deficiency: a systematic review and meta-analysis. Blood Cells Mol Dis, 2009. 42(3): p. 267–78.

2. Howes, R.E., et al., G6PD deficiency prevalence and estimates of affected populations in malaria endemic countries: a geostatistical model-based map. PLoS Med, 2012. 9(11): p. e1001339.

3. Cappellini, M.D. and G. Fiorelli, Glucose-6-phosphate dehydrogenase deficiency. Lancet, 2008. 371(9606): p. 64–74.

4. Peng, Q., et al., Large cohort screening of G6PD deficiency and the mutational spectrum in the Dongguan District in Southern China. PLoS One, 2015. 10(3): p. e0120683.

5. Minucci, A., et al., Glucose-6-phosphate dehydrogenase (G6PD) mutations database: review of the “old” and update of the new mutations. Blood Cells Mol Dis, 2012. 48(3): p. 154–65.

6. Perl, A., et al., Oxidative stress, inflammation and carcinogenesis are controlled through the pentose phosphate pathway by transaldolase. Trends Mol Med, 2011. 17(7): p. 395–403.

7. Beutler, E., Glucose-6-phosphate dehydrogenase deficiency: a historical perspective. Blood, 2008. 111(1): p. 16–24.

8. Luzzatto, L., C. Nannelli, and R. Notaro, Glucose-6-Phosphate Dehydrogenase Deficiency. Hematol Oncol Clin North Am, 2016. 30(2): p. 373–93.

9. Martini, G., et al., Structural analysis of the X-linked gene encoding human glucose 6-phosphate dehydrogenase. EMBO J, 1986. 5(8): p. 1849–55.

10. Takizawa, T., et al., Human glucose-6-phosphate dehydrogenase: primary structure and cDNA cloning. Proc Natl Acad Sci U S A, 1986. 83(12): p. 4157–61.

11. Kotaka, M., et al., Structural studies of glucose-6-phosphate and NADP+ binding to human glucose-6-phosphate dehydrogenase. Acta Crystallogr D Biol Crystallogr, 2005. 61 (Pt 5): p. 495–504.

12. Ho, H.Y., M.L. Cheng, and D.T. Chiu, Glucose-6-phosphate dehydrogenase--beyond the realm of red cell biology. Free Radic Res, 2014. 48(9): p. 1028–48.

13. Stanton, R.C., Glucose-6-phosphate dehydrogenase, NADPH, and cell survival. IUBMB Life, 2012. 64(5): p. 362–9.

14. Longo, L., et al., Maternally transmitted severe glucose 6-phosphate dehydrogenase deficiency is an embryonic lethal. EMBO J, 2002. 21(16): p. 4229–39.

15. Shu, L.P., et al., Ectopic expression of Hoxb4a in hemangioblasts promotes hematopoietic development in early embryogenesis of zebrafish. Clin Exp Pharmacol Physiol, 2015. 42(12): p. 1275–86.

16. Patrinostro, X., et al., A model of glucose-6-phosphate dehydrogenase deficiency in the zebrafish. Exp Hematol, 2013. 41(8): p. 697–710 e2.

17. Jiang, W., et al., Structure and function of glucose-6-phosphate dehydrogenase-deficient variants in Chinese population. Hum Genet, 2006. 119(5): p. 463–78.

18. Hejatko, J., et al., In situ hybridization technique for mRNA detection in whole mount Arabidopsis samples. Nat Protoc, 2006. 1(4): p. 1939–46.

19. Howes, R.E., et al., G6PD deficiency: global distribution, genetic variants and primaquine therapy. Adv Parasitol, 2013. 81: p. 133–201.

20. Rochford, R., et al., Humanized mouse model of glucose 6-phosphate dehydrogenase deficiency for in vivo assessment of hemolytic toxicity. Proc Natl Acad Sci U S A, 2013. 110(43): p. 17486–91.

21. Gomez-Gallego, F., A. Garrido-Pertierra, and J.M. Bautista, Structural defects underlying protein dysfunction in human glucose-6-phosphate dehydrogenase A(-) deficiency. J Biol Chem, 2000. 275(13): p. 9256–62.

22. Thermes, V., et al., I-SceI meganuclease mediates highly efficient transgenesis in fish. Mech Dev, 2002. 118 (1-2): p. 91–8.

23. Beutler, E. and M. Mitchell, Special modifications of the fluorescent screening method for glucose-6-phosphate dehydrogenase deficiency. Blood, 1968. 32(5): p. 816–8.

24. Mahmoud, A.A. and A.K. Nor El-Din, Glucose-6-Phosphate Dehydrogenase Activity and Protein Oxidative Modification in Patients with Type 2 Diabetes Mellitus. J Biomark, 2013. 2013: p. 430813.

25. Nadarajan, V., et al., Modification to reporting of qualitative fluorescent spot test results improves detection of glucose-6-phosphate dehydrogenase (G6PD)-deficient heterozygote female newborns. Int J Lab Hematol, 2011. 33(5): p. 463–70.

26. Nantakomol, D., et al., Evaluation of the phenotypic test and genetic analysis in the detection of glucose-6-phosphate dehydrogenase deficiency. Malar J, 2013. 12: p. 289.

27. Detrich, H.W., 3rd, et al., Intraembryonic hematopoietic cell migration during vertebrate development. Proc Natl Acad Sci U S A, 1995. 92(23): p. 10713–7.

28. De Domenico, I., et al., Zebrafish as a model for defining the functional impact of mammalian ferroportin mutations. Blood, 2007. 110(10): p. 3780–3.

29. Brownlie, A., et al., Positional cloning of the zebrafish sauternes gene: a model for congenital sideroblastic anaemia. Nat Genet, 1998. 20(3): p. 244–50.

