## Supplementary figures and images for "Establishment of a stable transgenic *g6pdM^1315-1443^* zebrafish line with glucose-6-phosphate dehydrogenase deficiency"

### Supplemental Figure 1

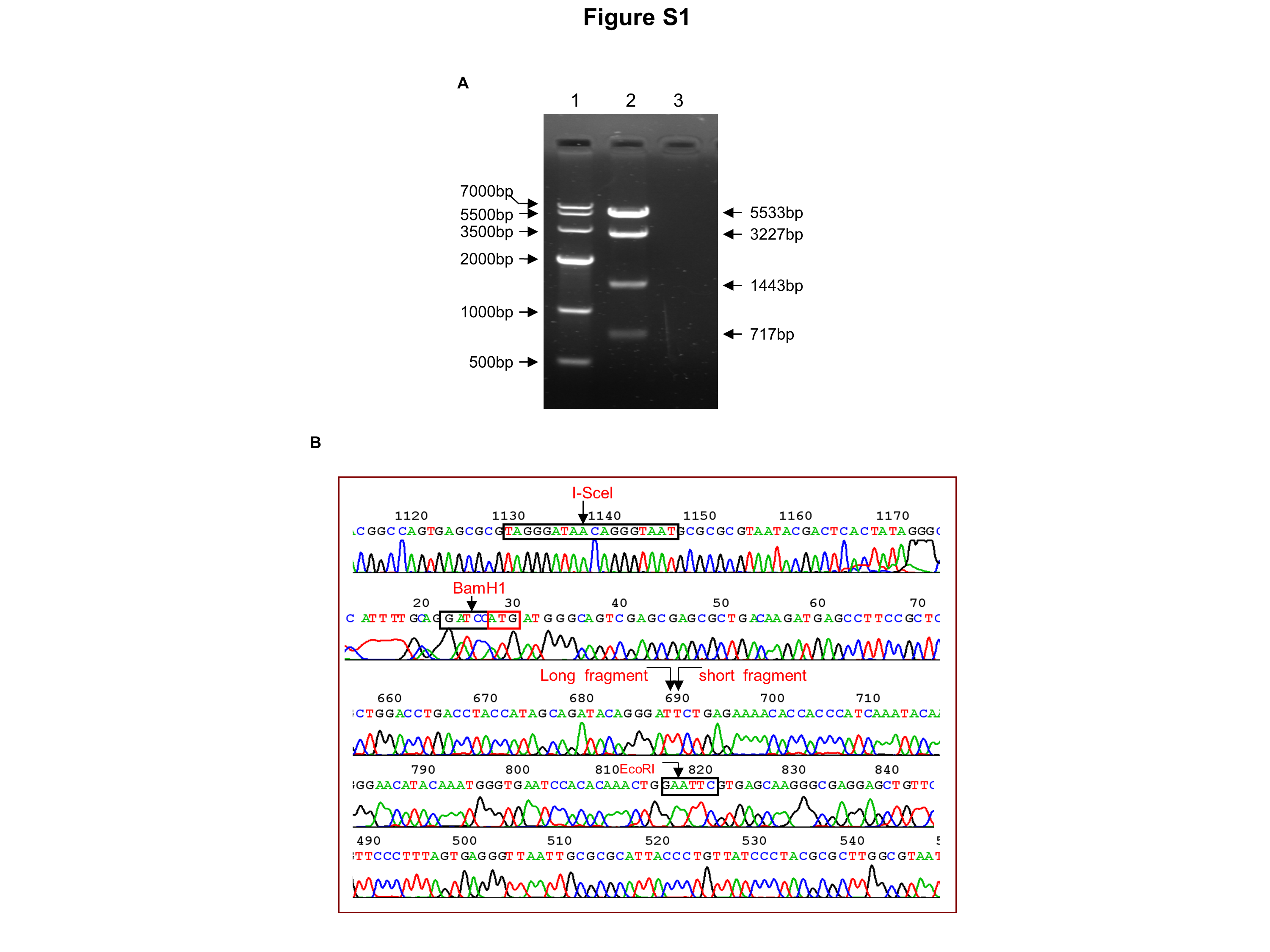

### Supplemental Figure 2

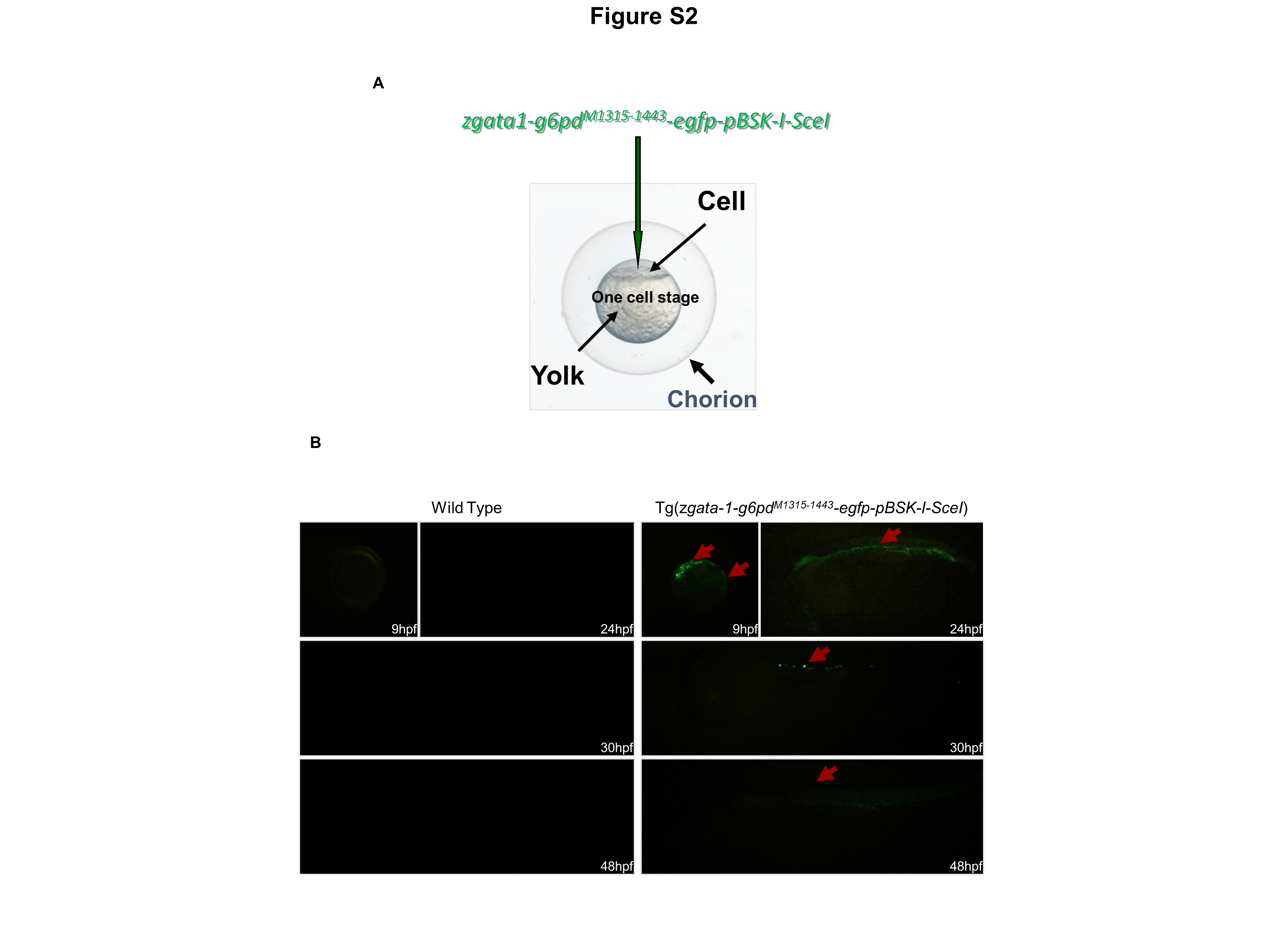

### Supplemental Figure 3

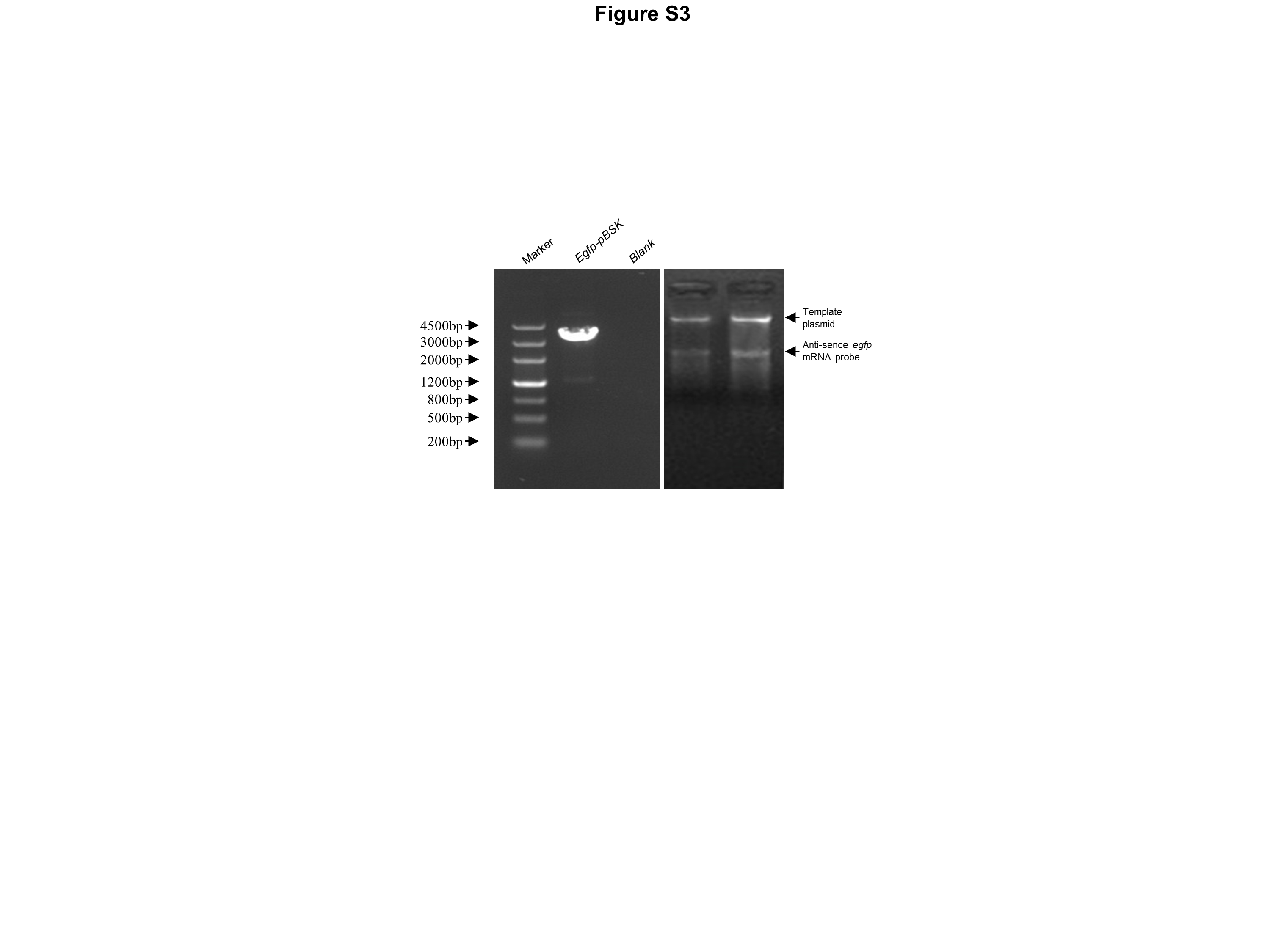
